# Spatio-temporal variability of soil nutrients and the responses of growth during growth stages of winter wheat in the north of China

**DOI:** 10.1101/398701

**Authors:** Su Bao-wei, Zhao Geng-xing, Dong Chao

## Abstract

Study on soil nutrient variability and its effect on the growth and development of crop under traditional tillage mode is the foundation to comprehensively implement the precision agriculture policy on the field scale and crop excellent management. In this paper, a winter wheat field of 28.5 hm2 under traditional cultivation model in Tianzhuang town of Huantai county was selected as the research area. Sampling by mesh point method (60×60m), the variation characteristics of soil available nitrogen (AN), available phosphorus (AP) and available potassium (AK) in the before sowing, reviving stage, jointing stage, filling stage of winter wheat were analyzed by the method of geostatistics and GIS. At the same time, Pearson correlation analysis was used to study the response of wheat growth and development to soil nutrient variation. As the growth period goes by, 1) each nutrient showed low-high-low and moderate variability. AN and AK had the highest content at reviving stage, while AP at jointing stage, as turning point. The order of variability of each nutrient was AN>AP>AK. 2) The difference of nutrient variation first increased and then decreased, and showed medium to strong spatial correlation.

Three nutrients in the before sowing stage were strong spatial correlation, and the reviving stage was medium spatial correlation, to the jointing and filling stages, AN was medium spatial correlation, AP and AK were strong spatial correlation. The spatial correlation of each nutrients was the weakest in the reviving stage, and AN was the strongest in the before sowing stage, while AP and AK were strongest in the jointing stage. The spatial correlation of each soil nutrients decreased from before sowing to reviving stage, jointing stage to filling stage, and the spatial correlation increased from reviving stage to jointing stage. 3) Soil nutrient content first increased and then decreased, and the grade of nutrient gradually decreased. 4) The correlation between soil nutrients and wheat growth was gradually increasing. AN had the highest correlation with wheat growth, followed by AK and AP lowest. The effect of soil nutrients on the growth of wheat at the reviving stage was higher than that of the current nutrient, and the growth of wheat at jointing stage was mainly influenced by the current nutrient, while the growth of wheat at the filling period was both influenced by the nutrient content of the last and the current period; the date to supplement fertilizer should be postponed properly. In this study, the soil nutrient dynamics and its influence on the growth of wheat during the winter wheat growth period under the traditional field model were well understood, which could provide a theoretical basis for the precision management of soil nutrients in the northern winter wheat area with relatively uniform planting environment and cultivation management.

## Introduction

The research on soil and crop under field conditions is a key step in precision agriculture from theory to practice. Soil, continuous variant of time and space, is the carrier and nutrient source of plant growth [1-3]. It is basis for precision management and fertilization of farmland to grasp temporal and spatial variability of soil nutrients and the crop responses in different growth stages in time [4-7].

Soil nutrient variability has long been a research hotspot of scholars all over the world [8-10], and many experiments have been carried out on various nutrients and different scales. For example, Duraisamy et al. [11] studies the spatial heterogeneity of soil PH, organic carbon and available nutrients in the typical dry areas of India. Rosemary et al. [12] studied the spatial structure of topsoil under multi-scale and intensity. Paz et al. [13] explained the spatial dependence of soil nutrients and trace elements even in very small areas (about 1.8 hectares). Yang QY et al. [14]compared the scale effects of soil AP and AK on spatial variability under different sampling scales. Blanchet et al. [15]analyzed the effects of different land use patterns, soil types, topography and other factors on spatial variability of soil K. However, in general, most of the studies focus on the spatial variability of soil nutrients, less consideration on the temporal variability and effects to crop growth. Thus, the spatio-temporal variability of soil nutrients in different growth stages of crops remains to be strengthened [16-18].

Santillano-Cázares et al. [19] showed that the temporal variability of soil nutrients was much greater than its spatial variability. Qiao JF et al. [20] studied the temporal and spatial variation characteristics of soil in wheat fields with different drip irrigation years. However, these studies are still limited to the analysis of soil nutrient itself, and the research of soil nutrient variability combined with crop development is still very scant. Plant growth depends on the absorption of soil nutrients, so there should also be spatio-temporal variability of plant growth status caused by soil nutrient variability [21-24]. The analysis of the variability of soil nutrient and its correlation with the crop growth will help to clarify the spatio-temporal variability of the variables in the farmland system and have positive productive significance [25-27]. Meanwhile, we found that previous studies in this area mostly used the method of field experiment design. Although this method can obtain better test results by controlling the test conditions, the conclusion has great limitations in the practical application of agricultural extension because it is different from the real field conditions.

Based on this, the variation characteristics of soil nutrients and their effects on wheat growth were analyzed by investigating the traditional cultivation management model of the main winter wheat production areas in China. So as to find out the drawbacks of this model and provide scientific basis for fertilization in different growth stages of wheat.

## Materials and methods

### Study area

The study was conducted in Huantai County with an area of 28.5 hm^2^, located in the North Central of Shandong province, China (37° 1.98’ ∼ 37°2.2S’N, 117°59.7’ ∼ 11 S°0.12’E). It is located in the Huanghuai winter wheat region, most important wheat-producing area in China. The region is classified as having a continental monsoon climate of warm temperate zone. The climate is mild and the four seasons are clear. The average annual air temperature is 12.5°C and frost-free period is 197 days. The annual average precipitation is 587mm, mainly in 6∼9 months. The terrain is flat with an elevation of 5.7 to 6.8m. The main soil types are cinnamon soil and the winter wheat variety is Jimai 17, and the whole process is the traditional cultivation management mode of the farmers. It is sowed in October 13, 2016 and harvested in June 10, 2017, with a total growth period of 260 days. Among them, basal compound fertilizer [N + P_2_O_5_ + K_2_O < 45% (15-15-15)] was applied before sowing on October 12, 2016, the Dosage is 360kg/hm^2^ and urea [CO (NH_2_)_2_] was additional applied on March 5, 2017, the Dosage is 300kg/hm^2^.

### Soil sampling and field survey

First, we collected the basic maps of the land use status map, soil map and topographic map of Huantai County, and then obtained the boundary of the study area with the help of ArcGIS and Google earth, finally, a total of 81 sampling points were set up according to the standard of uniform grid layout according to 60m×60m (fig. 1). The coordinates of the sampling points were transformed and input into the Trimble Geo 7x series GPS, its positioning accuracy can reach centimeter level. Field surveys were conducted on October 8, 2016 (before sowing) and March 10, 2017 (reviving stage), April 10 (jointing stage) and May 15, 2017 (filling stage).

**Fig.1.**
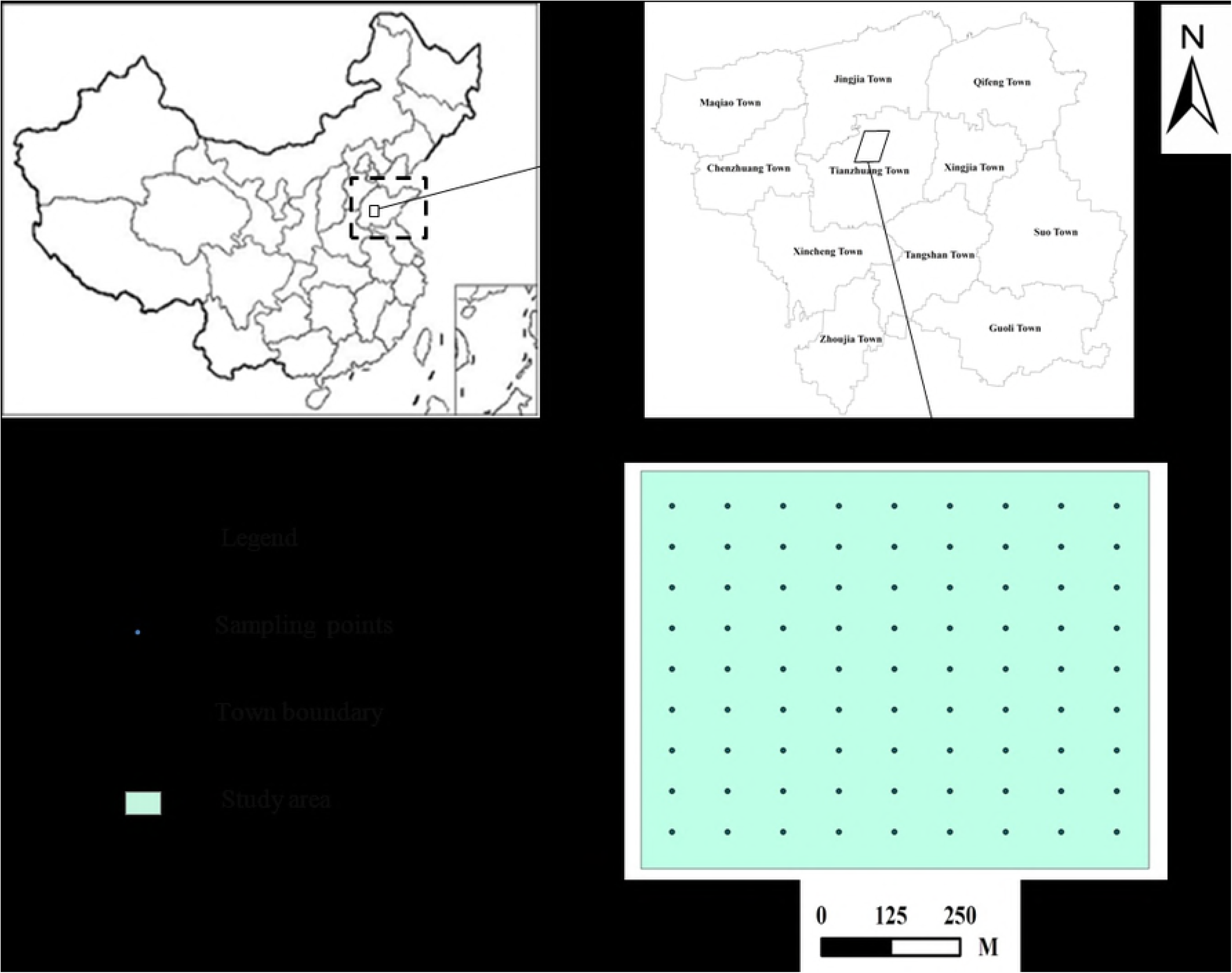
Location of die study area and distribution of soil sampling sites

Soil samples were collected by five-point cross-sampling method with a sampling depth of 0-20 cm. Multi-point soil samples were air-dried, crushed, sieved, and analysed for soil AN, AP, AK through the method of agro-chemistry (Nanjing Agriculture College. 1980). The chlorophyll content (SPAD) of leaves of wheat was measured by SPAD-502 chlorophyll content analyzer. 5 wheat plants were randomly selected at each observation point, and 3 times per plant were measured, average value of the 5 wheat was calculated as the SPAD value of this sampling point. The leaf area index (LAI) of wheat was measured by LAI-2200C plant canopy analyzer, each sampling point was measured by two methods: cross the ridge and parallel to the ridge, each method was collected for 3 times. Finally, the mean value was obtained as the LAI value of this point.

## Data processing and analysis methods

### Data processing

The data were screened with SPSS 21.0 software. Interval method was used to select abnormal values. The interval is [μ-3s, μ+3s]. μ was the mean value of sample data, s was the standard deviation, and the outliers outside the interval were replaced by the normal maximum and minimum value.

Data normal distribution was checked with the Kolmogorov-Smirnov method. The data that did not conform to the normal distribution should be converted. It showed that the distribution of sample data was not significantly different from normal distribution when sig> 0.05.

### Analysis methods

The eigenvalues of the classical statistical methods include minimum, maximum, mean, standard deviation and coefficient of variation (CV) et al, which can be used to describe the basic physical characteristics of soil nutrients. Among them, the coefficient of variation is a parameter that can be used to compare the degree of data discretization under different measurement scales, according to its classification level, <10% indicates weakly variability, 10% to 100% indicates medium variability, and >100% is strongly variability. Single factor analysis of variance-least significant difference (LSD) method in SPSS software was used to analyze the significant difference of soil nutrients in different growth stages.

The spatio-temporal variability of soil nutrients was mainly analyzed by semi-variogram, Moran ’ s I and fractal dimension (D) was combined to determine its spatial autocorrelation and agglomeration degree at the same time. Formula for the Moran ’ s I was as follows:

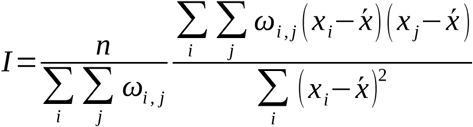

The value of Moran’ s I was between −1 and 1, when the adjacent region has a similar attribute value, the value of Moran ’ s I was positive; When the adjacent area was different value, it is negative; when the attribute value is pure random arrangement, it tends to 0. First, the data of AN, AP and AK of each growth period were input to Gs+ software respectively, and the distribution point maps were obtained by semi-variogram, then the point maps were fitted with spherical, exponential, Gauss and linear models respectively. Finally, the best model and its corresponding parameter index were obtained. Among them, C_0_/ (C_0_+C) was an important parameter to test the degrees of variation in the system variables. When C_0_/ (C_o_+C) <25%, it indicated that the system has strong spatial correlation; when it was between 25% and 75%, this indicated that the system has moderate spatial correlation; when >75%, this indicated that the system has weak spatial correlation.

The spatial distribution map of soil AN, AP and AK in different growth stages was plotted with the method of Kriging interpolation. The classification standard of soil nutrient of the interpolation map was formulated according to China’s second national soil survey. Among them, AN is divided into 3 categories: high (**I, II**), medium **(III, IV)** and low (**V, VI)** 6 grades, AP and AK divided into 3 categories: high (**I, II),** medium (III, **IV, V**) and low **(VI,** VII) 7 grades. On the basis of nutrient interpolation map, the professional viewpoints of landscape ecology were referred. 2 indexes of landscape level: patch number (NP), patch density (PD) and 1 index of type level: the percentage of landscape area (PLAND). A total of 3 indexes were selected to analyze the spatial dynamic characteristics of soil nutrient variation quantitatively.

The values of SPAD, LAI at different growth stages of wheat and soil AN, AP, AK were corresponded to point by point. The correlation coefficient was calculated by using Pearson correlation analysis to analyze the response of winter wheat growth to soil nutrient change.

## Results and analysis

### The descriptive statistical analysis of soil nutrients in different growth stages of Winter Wheat

There were the statistical characteristics of soil nutrients in different growth stages of winter wheat (Table 1). The content of AN ranged from 50.82 to 91.24 mg/kg and the coefficient of variation ranged from 51.50% to 56.47% from before sowing to filling stage. The content of AN in before sowing and reviving stage was significantly higher than that in jointing and filling stage. Reviving stage was the turning point of the change of soil AN content. The content increased slightly before this period, and then decreased significantly after that. There was no significant difference in AN content between before sowing and reviving stages, jointing and filling stages, but there was significant difference in AN content between the first two and the last two growth stages. The content of AP ranged from 35.70 to 64.37 mg/kg and the coefficient of variation ranged from 34.56% to 59.14% from before sowing to filling stage. The content of AP was low before sowing, and was significantly different from that of the other three growth stages. At the reviving stage, content of AP increased significantly and increased slightly at jointing stage, then decreased slightly at filling stage. The coefficient of variation of AP was moderate in each growth stages, but the coefficient of variation before sowing was obviously higher, followed by jointing stage, and the lowest at reviving stage and filling stage. The content of AK ranged from 285.19 to 377.13 mg/kg and the coefficient of variation ranged from 16.84% to 28.06% from before sowing to filling stage. The content of AK in the reviving period was significantly higher than that of other periods. The content of it increased significantly from before sowing to reviving stage, but decreased significantly at jointing stage. The coefficient of variation of AK was moderate in each growth stages, slightly higher before sowing, lower at jointing stage, middle at reviving stage and filling stage, but the difference was not obvious.

**Table 1:**
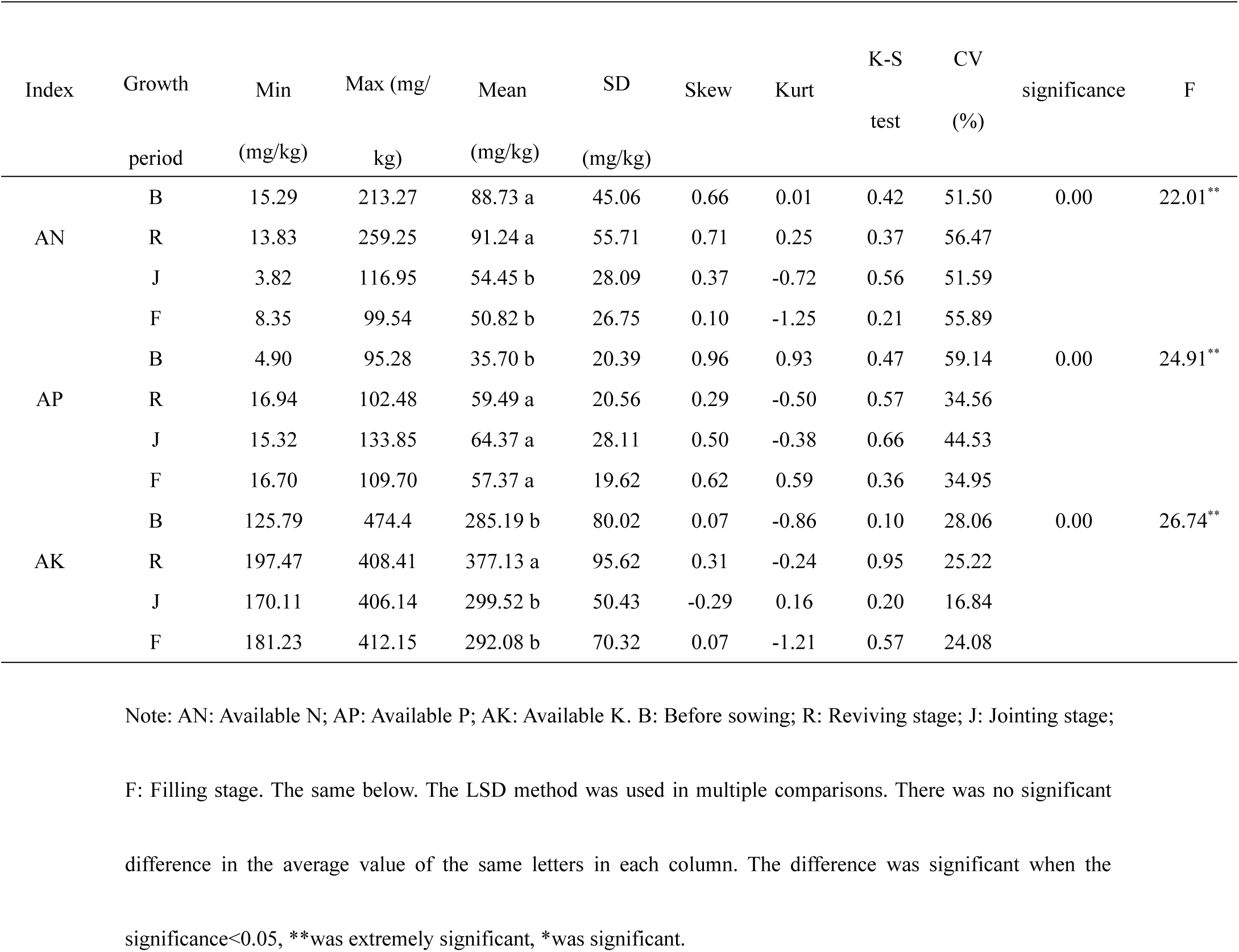
Statistical characteristic of soil nutrients during the growth period of winter wheat

In general, the content of soil nutrients showed the characteristics of low-high-low and moderate variability at different growth stages. The content of AN, AK were the highest in the period of reviving stage and in the jointing stage of AP, according to grading standard of China’s second national soil survey, the AN content in the study area was relatively low, while AP and AK were abundant. The order of the degree of variation of each nutrient was AN>AP>AK. The results of kolmogorov-Smirnov test showed that under the 5% test standard, AN, AP and AK of each growth period obeyed normal distribution after logarithmic transformation, which could meet the requirements of Moran’s I and Kriging interpolation.

### The variation characteristics of soil nutrients in different growth stages of Winter Wheat

There were the semivariogram models of soil nutrients in different growth stages of winter wheat (Figure 1). It could be seen that the best fitting models of the three nutrients before sowing were all exponential models, indicating that each nutrient has a similar trend of spatial variation, and the combination of the variation characteristics of soil nutrients was the simplest in this growth period. The best fitting model of AN was exponential model, while AP and AK were spherical model at reviving stage. It showed that the variation trend of AP and AK was similar, but different from that of AN in this period. To jointing stage, the combination of soil nutrient variation characteristics was the most complicated, contrary to that before sowing. The best fitting models of AN, AP and AK were spherical, exponential and Gauss models in turn. To filling stage, the best fitting models of AN and AP were Gaussian model and AK was exponential model, which showed that the spatial variation trends of AN and AP were similar, but different from AK, the soil nutrient variation characteristics of this growth period and reviving stage were similar in complexity, between the before sowing and jointing stages.

In general, the semi-variogram of the three nutrients were all the same model before sowing, to the two models at reviving stage, to the three models at jointing stage, and then to two models in the filling period. The variation characteristics were greatest at jointing stage, and the difference was higher and then decreased. The R^2^ range of the best fitting model of each nutrient was 0.904 to 0.997, and the range of RSS was 3.36×l0^−8^ to 6.93×10^−5^, which indicated that the selected model of semi-variogram was better and could reflect the spatial structure characteristics and variation trend of soil nutrients during the growth period well.

There were the semivariogram models and parameters for soil nutrients in different growth stages of winter wheat (Table 2). It could be seen that the C_0_ of AN, AP and AK in the four growth stages were all close to 0, indicated that the variation caused by test and sampling errors was very small. The C_0_/ (C_0_+C) of AN was 10.29% before sowing, with strong spatial correlation, and it ranging from 27.09% to 38.67% from reviving stage to filling stage, with moderate spatial correlation. The C_0_/ (C_0_+C) of AP before sowing, jointing and filling stages were 8.65%, 8.35% and 15.89%, respectively, with strong spatial correlation, and 34.35% at reviving stage, moderate spatial correlation. The C_0_/ (C_0_+C) of AK before sowing, jointing and filling stages were 12.5%, 10.77 and 13.82%, respectively, with strong spatial correlation, and 26.09% at reviving stage, moderate spatial correlation. Overall, the three nutrients showed moderate to strong spatial correlation with the development of growth period, and experienced a process of weakening-strengthening-weakening. From the angle of growth period, these three nutrients were strong spatial correlation before sowing, and the reviving period was moderate spatial correlation, to the jointing and filling stages, AN was moderate spatial correlation, and AP and AK were strong spatial correlation. From the angle of nutrient, these three nutrients were the weakest spatial autocorrelation at reviving stage, the before sowing of AN was the strongest, while the AP and AK were the strongest at jointing stage.

**Table 2:** Semivariogram theoretical models and parameters of soil nutrients during the growth period of winter wheat

| Growth<br>period | Index | model | $C_0$ | $C_0+C$ | $C_0/(C_0+C)$<br>(%) | A<br>(m) | $R^2$ | RSS | D | $I$ |
| --- | --- | --- | --- | --- | --- | --- | --- | --- | --- | --- |
| Before<br>sowing | AN | E | 0.0060 | 0.0583 | 10.29 | 245.7 | 0.941 | $1.68 \times 10^{-5}$ | 1.857 | 0.305 |
| | AP | E | 0.0073 | 0.0844 | 8.65 | 164.7 | 0.993 | $2.13 \times 10^{-6}$ | 1.897 | 0.226 |
| | AK | E | 0.0024 | 0.0192 | 12.5 | 235.2 | 0.976 | $7.79 \times 10^{-7}$ | 1.817 | 0.320 |
| Reviving<br>stage | AN | E | 0.0408 | 0.1055 | 38.67 | 179.8 | 0.969 | $2.10 \times 10^{-5}$ | 1.887 | 0.282 |
| | AP | S | 0.0090 | 0.0262 | 34.35 | 142.5 | 0.920 | $3.08 \times 10^{-6}$ | 1.925 | 0.209 |
| | AK | S | 0.0036 | 0.0138 | 26.09 | 235.8 | 0.997 | $8.70 \times 10^{-8}$ | 1.914 | 0.153 |
| Jointing<br>stage | AN | S | 0.0233 | 0.0860 | 27.09 | 146.8 | 0.913 | $4.92 \times 10^{-5}$ | 1.855 | 0.280 |
| | AP | E | 0.0035 | 0.0419 | 8.35 | 122.7 | 0.907 | $5.49 \times 10^{-6}$ | 1.900 | 0.212 |
| | AK | G | 0.0007 | 0.0065 | 10.77 | 108.1 | 0.982 | $3.36 \times 10^{-8}$ | 1.853 | 0.260 |
| Filling<br>stage | AN | G | 0.0333 | 0.0916 | 36.35 | 127.5 | 0.904 | $6.93 \times 10^{-5}$ | 1.890 | 0.270 |
| | AP | G | 0.0048 | 0.0302 | 15.89 | 88.2 | 0.938 | $2.20 \times 10^{-6}$ | 1.941 | 0.177 |
| | AK | E | 0.0017 | 0.0123 | 13.82 | 117.9 | 0.929 | $2.72 \times 10^{-7}$ | 1.934 | 0.125 |

Note: E: Exponential model; S: Spherical model; G: Gaussian model. D: The fractal dimensions. $I$ : Moran 's I.

The fractal dimensions of the three nutrients were from 1.817 ∼ 1.897, 1.887 ∼ 1.925, 1.853 ∼ 1.900 and 1.890 ∼ 1.941 from before sowing to filling stage, indicating that the soil nutrients in each growth period had good fractal characteristics and structure. Its value showed a process of firstly increasing then decreasing and then increasing from before sowing. It indicated that the structure of soil nutrient became more complex and the spatial correlation was weakened from before sowing to reviving stage, jointing stage to filling stage; while the structure of nutrients tended to be consistent and spatial correlation increased from reviving to jointing stage.

There were the parameters of Moran’s I for soil nutrients in different growth stages of winter wheat (Table 2). It could be seen that Moran’s I of the three nutrients range from 0.226 ∼ 0.320, 0.153 ∼ 0.282. 0.212 ∼ 0,280 and 0.125 ∼ 0.270 from the period of before sowing to filling stage, and the values were between 0 and 1. It indicated that there was a regular concentration distribution of the three nutrients at different growth stages. The value of Moran’s I showed a process of firstly decreasing then increasing and then decreasing from before sowing. The results showed that soil nutrient structure was gradually dispersed and the spatial correlation decreased from before sowing to reviving stage, jointing stage to filling stage; while the structure of nutrients tends to gather and spatial correlation increased from reviving stage to filling stage.

In general, as the growth period goes on: (1) each nutrient showed moderate to strong spatial correlation. (2) The three nutrients were strong spatial correlation before sowing; and the reviving period was moderate spatial correlation; to the jointing and filling stages, AN was moderate spatial correlation, and AP and AK were strong spatial correlation. (3) The three nutrients were the weakest spatial autocorrelation at reviving stage; the before sowing of AN was the strongest, while the AP and AK were the strongest at jointing stage. (4) The autocorrelation of soil nutrient decreased from before sowing to reviving stage, jointing stage to filling stage; and increased from reviving stage to jointing stage.

### The spatial distribution characteristics of soil nutrients in different growth stages of Winter Wheat

There were the spatial and temporal distribution maps of soil nutrients in different growth stages of winter wheat (Figure 4). It could be seen that the distribution of AN before sowing was strip shaped; and the content from northwest to southeast was obviously higher than that of other regions. To reviving stage, the concentration of AN distribution declined slightly; and the content of the area was opposite to that before sowing, area with high content was concentrated in the direction of northeast to southwest. To jointing stage, AN showed a lump cluster distribution; and the content of it decreased significantly, mainly concentrated in 60∼90 mg/kg. To filling stage, the continuity of the distribution continued to increase; and the content continued to decline, its value was lower than 60 mg/kg. The distribution of AP before sowing was dispersed, and there was no clear center of high, low content. To reviving stage, the distribution of AP showed strip shaped and the content increased significantly. To jointing stage, the boundary of high and low regional were obvious, the overall trend was high in the east and low in the west and the content of the study area was higher, mainly concentrated in > 60 mg/kg level. To filling stage, AP showed a lump cluster distribution and the area of high nutrient content was reduced. The distribution of AK before sowing showed a lump cluster distribution and the high content area was concentrated in the northeast. To reviving stage, the distribution of AK showed strip shaped and the area of high content became larger. To jointing stage, the continuity of AK distribution decreased significantly, and the area with high content decreased rapidly. To filling stage, the distribution characteristics of AK were similar to that at jointing stage; the content decreased further and the area with low content increased.

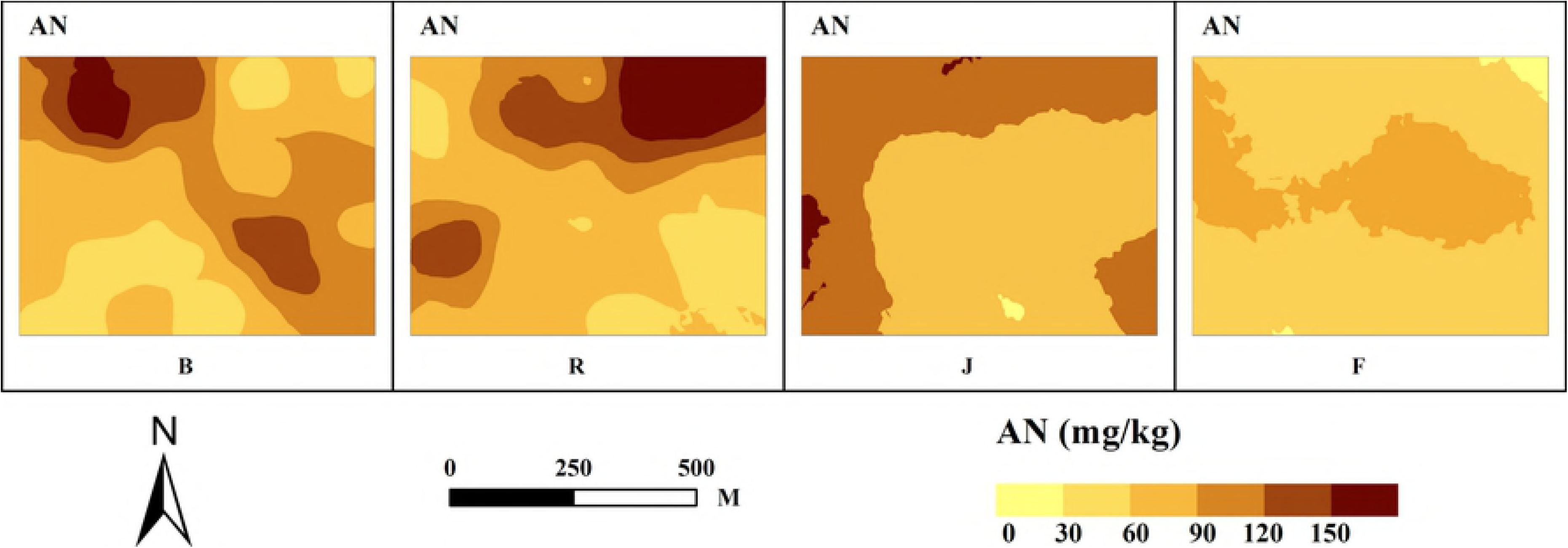

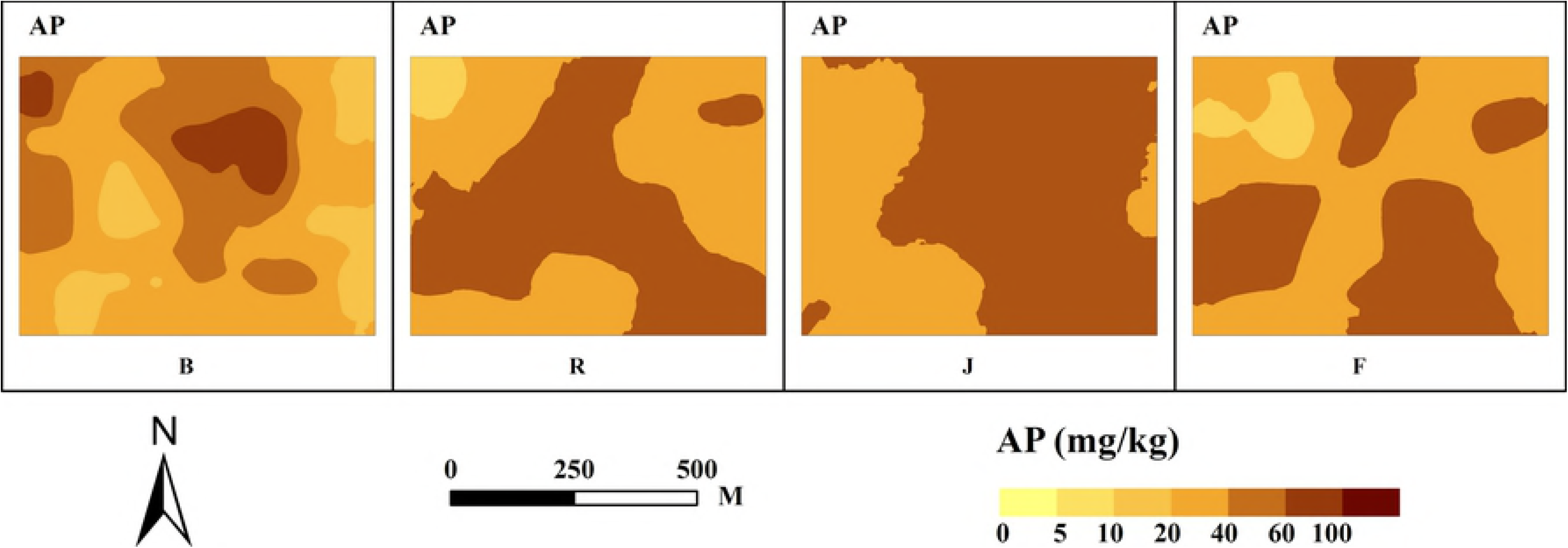

**Fig.4.**
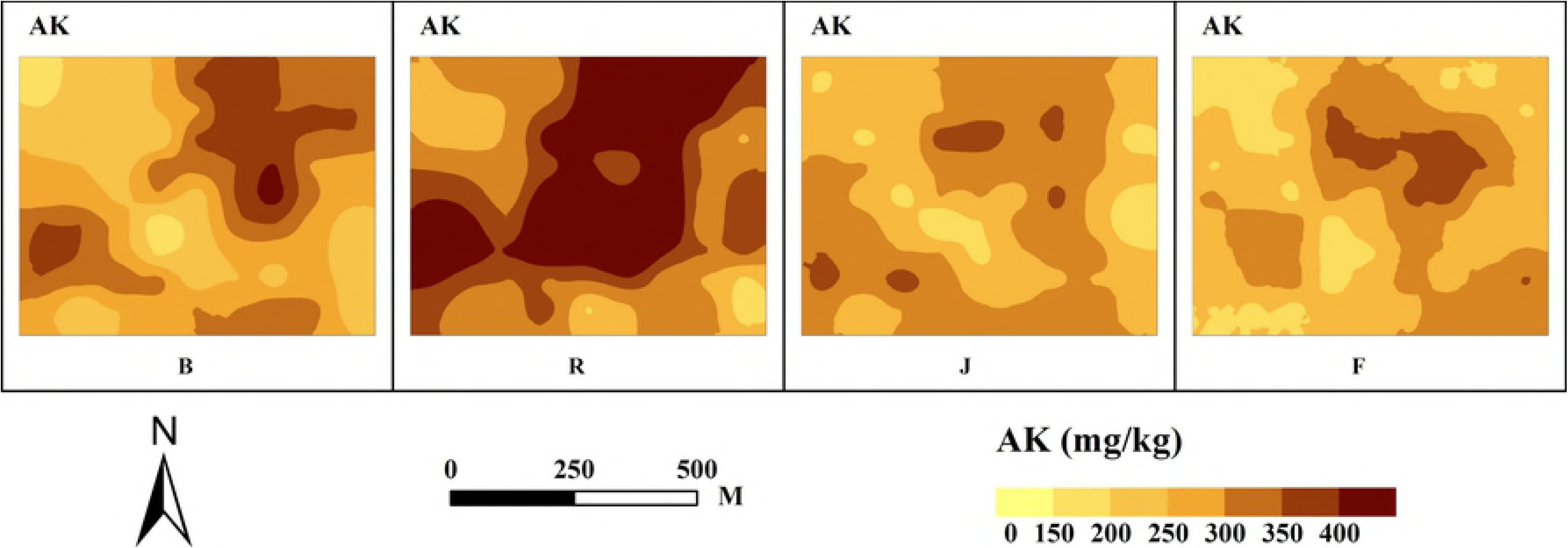
Spatial distribution of soil nutrients during the growth period of Wheat

There were the landscape pattern indices for the interpolation map of soil nutrients in different growth stages of winter wheat (Table 3-5). The distribution of AN was consistent at landscape level during different stages. Patch number and patch density showed the characteristics of increased-decreased-decreased. From the angle of type level, the proportion of high grade (I and II) was 20.39%, 28.32%, 0, 0, the middle grade (III and IV) was 55.79%, 50.56%, 49.33%, 27.6% and the low grade (V and VI) was 23.82%, 21.11%, 50.66%, 72.4% from before sowing to filling stage. This showed that with the development of the growth period, the proportion of the high and low levels of AN was converted, the proportion of the high content gradually decreases after the reviving stage, while the proportion of the low grade started to increase. The distribution of AP was consistent at landscape level during different stages. Patch number and patch density showed the characteristics of decreased-decreased-increased. From the angle of type level, the proportion of high grade (I and II) was 10.18%, 55.23%, 67.81%, 48.77%, the middle grade (III, IV, V) was 88.28%, 44.77%, 32.17%, 51.23% and the low grade (VI and VII) was 1.53%, 0, 0, 0 from before sowing to filling stage. This showed that the dominant areas converted from high content to middle content. The distribution of AK was consistent at landscape level during different stages, showed a characteristic of continuous increase. From the angle of type level, the proportion of high grade (I and II) was 18.13%, 55.59%, 6.38%, 6.34%, the middle grade (III, IV, V) was 76.4%, 44.41%, 93.62%, 92.56% and the low grade (VI and VII) was 5.46%, 0, 0, 1.1% from before sowing to filling stage. This showed that with the development of the growth period, change rule of the AK at type level was same as that of AN in general, and the proportion of high grade decreased after reviving stage. Generally, in the whole growth period of winter wheat, the contents of soil nutrients were firstly increased and then gradually decreased, while the number of nutrient grades, especially the number of high content grades, gradually decreased.

**Table 3:** Variation of AN landscape indices at landscape and class level during the growth period of winter wheat

| Growth period | Index | Class I | Class II | Class III | ClassIV | Class V | ClassVI |
| --- | --- | --- | --- | --- | --- | --- | --- |
| Before sowing | NP | 1 | 2 | 1 | 5 | 3 | 1 |
|  | PD (/hm<sup>2</sup>) | 0.0286 | 0.0571 | 0.0286 | 0.1429 | 0.0857 | 0.0286 |
|  | PLAND | 4.52% | 15.87% | 18.56% | 37.23% | 22.87% | 0.95% |
| Reviving stage | NP | 1 | 2 | 2 | 6 | 3 |  |
|  | PD (/hm<sup>2</sup>) | 0.0286 | 0.0571 | 0.0571 | 0.1714 | 0.0857 |  |
|  | PLAND | 14.38% | 13.94% | 16.55% | 34.01% | 21.11% |  |
| Jointing stage | NP |  |  | 5 | 2 | 1 | 1 |
|  | PD (/hm<sup>2</sup>) |  |  | 0.1429 | 0.0571 | 0.0286 | 0.0286 |
|  | PLAND |  |  | 1.76% | 47.57% | 50.23% | 0.43% |
| Filling stage | NP |  |  |  | 2 | 4 | 2 |
|  | PD (/hm<sup>2</sup>) |  |  |  | 0.0571 | 0.1143 | 0.0571 |
|  | PLAND |  |  |  | 27.60% | 71.27% | 1.13% |

Note: NP: patch number; PD: patch density; PLAND: percentage composition of landscape. The same below.

**Table 4:** Variation of AP landscape indices at landscape and class level during the growth period of winter wheat

307Table 4 Variation of AP landscape indices at landscape and class level during the growth period of winter wheat
| Growth<br>period | Index | Class I | Class II | Class III | ClassIV | Class V | ClassVI | ClassVII |
| --- | --- | --- | --- | --- | --- | --- | --- | --- |
| Before<br>sowing | NP |  | 2 | 3 | 2 | 5 | 1 | 1 |
|  | PD (/hm<sup>2</sup>) |  | 0.0571 | 0.0857 | 0.0571 | 0.1429 | 0.0286 | 0.0286 |
|  | PLAND |  | 10.18% | 31.96% | 39.11% | 17.21% | 0.81% | 0.72% |
| Reviving<br>stage | NP | 1 | 3 | 3 | 1 |  |  |  |
|  | PD (/hm<sup>2</sup>) | 0.0286 | 0.0857 | 0.0857 | 0.0286 |  |  |  |
|  | PLAND | 0.92% | 54.31% | 41.38% | 3.39% |  |  |  |
| Jointing<br>stage | NP |  | 2 | 3 |  |  |  |  |
|  | PD (/hm<sup>2</sup>) |  | 0.0571 | 0.0857 |  |  |  |  |
|  | PLAND |  | 67.81% | 32.17% |  |  |  |  |
| Filling<br>stage | NP |  | 4 | 1 | 1 |  |  |  |
|  | PD (/hm<sup>2</sup>) |  | 0.1143 | 0.0286 | 0.0286 |  |  |  |
|  | PLAND |  | 48.77% | 45.35% | 5.88% |  |  |  |

**Table 5:** Variation of AK landscape indices at landscape and class level during the growth period of winter wheat

308Table 5 Variation of AK landscape indices at landscape and class level during the growth period of winter wheat
| Growth<br>period | Index | Class I | Class II | Class III | ClassIV | Class V | ClassVI | ClassVII |
| --- | --- | --- | --- | --- | --- | --- | --- | --- |
| Before<br>sowing | NP | 1 | 2 | 3 | 1 | 4 | 2 | 1 |
|  | PD (/hm<sup>2</sup>) | 0.0286 | 0.0571 | 0.0857 | 0.0286 | 0.1143 | 0.0571 | 0.0286 |
|  | PLAND | 1.83% | 16.30% | 21.99% | 26.13% | 28.28% | 3.95% | 1.51% |
| Reviving<br>stage | NP | 2 | 3 | 3 | 4 | 2 |  |  |
|  | PD (/hm<sup>2</sup>) | 0.0571 | 0.0857 | 0.0857 | 0.1143 | 0.0571 |  |  |
|  | PLAND | 33.87% | 21.72% | 30.19% | 12.86% | 1.36% |  |  |
| Jointing<br>stage | NP | 1 | 5 | 1 | 3 | 6 |  |  |
|  | PD (/hm<sup>2</sup>) | 0.0286 | 0.1429 | 0.0286 | 0.0857 | 0.1714 |  |  |
|  | PLAND | 1.24% | 5.14% | 55.85% | 31.46% | 6.31% |  |  |
| Filling<br>stage | NP |  | 2 | 2 | 7 | 9 | 1 |  |
|  | PD (/hm<sup>2</sup>) |  | 0.0571 | 0.0571 | 0.2 | 0.2571 | 0.0286 |  |
|  | PLAND |  | 6.34% | 31.16% | 47.27% | 14.13% | 1.10% |  |

In general, with the development of the growth period, (1) the spatial distribution continuity of soil AN increased continuously, and the high - middle - low content level was characterized the trend of vanishing - decreasing - increasing. (2) AP took the jointing period as the turning point, the spatial distribution continuity was increased before the growth period, and the proportion of high level was gradually increased; while the continuity of spatial distribution decreases, the dominant areas converted from high content to middle content after the growth period. (3) The spatial distribution continuity of AK continued to decline, and the high content level was decreased significantly after reviving stage.

### Correlation between soil nutrient and the index of wheat growth

There were the correlation coefficients between soil nutrient and the index of wheat growth (Table 6). It could be seen that there was a good correlation between AN, AP, AK and SPAD, LAI of wheat. Among them, SPAD, LAI and AN reached a significant positive correlation at each growth stage. There was no significant correlation between SPAD and AP at any growth stages. There was a significant correlation between LAI and AP at filling stage, but there was no correlation during the period of reviving stage to jointing stage. There was a significant correlation between SPAD and AK at filling stage, but there was no correlation during the period of reviving stage to jointing stage. There was no correlation between LAI and AK at filling stage, but it reached significant correlation during the period of reviving stage to jointing stage.

**Table 6:**
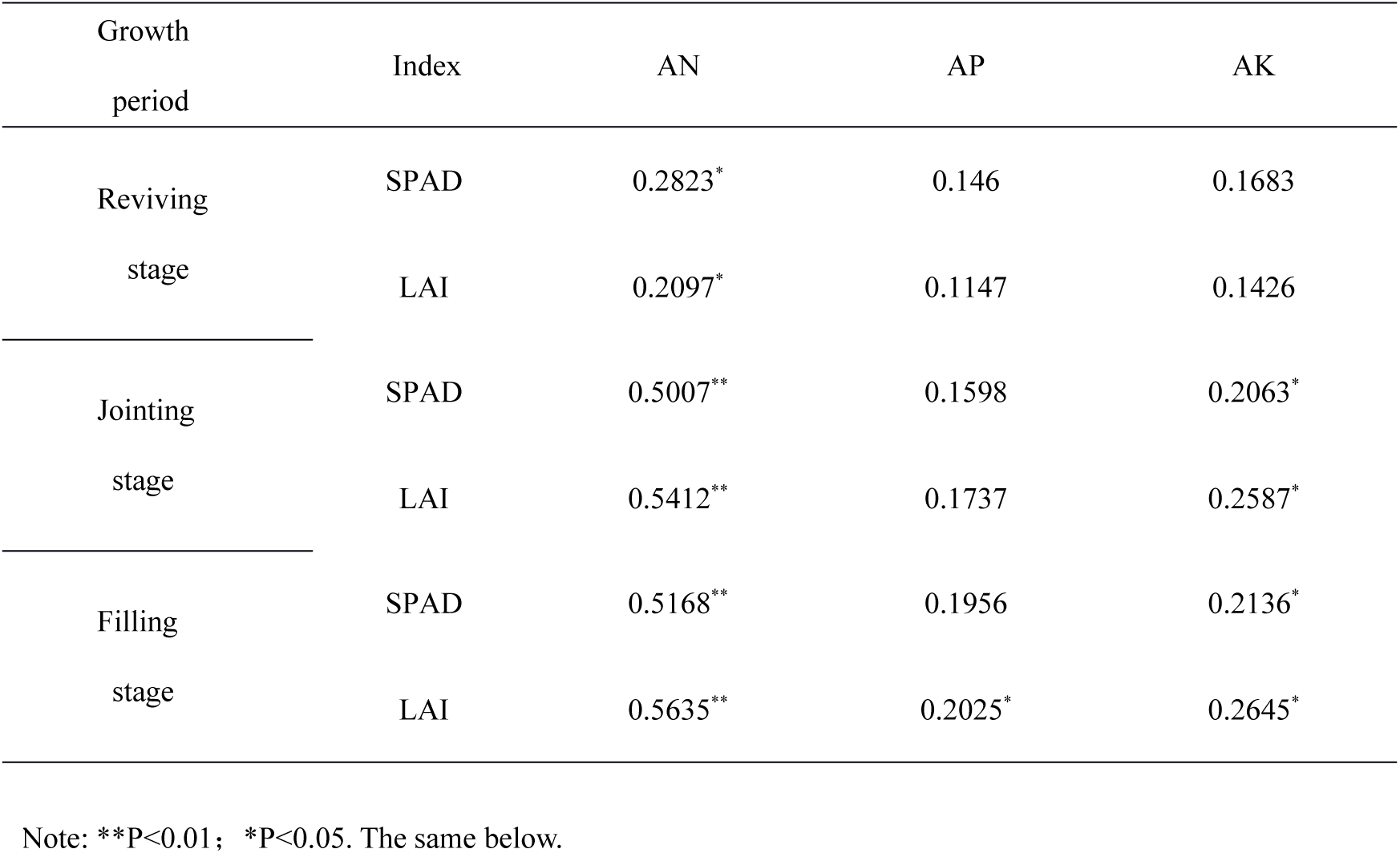
Correlation coefficients between soil nutrients and wheat fertility index

| Growth period | Index | AN | AP | AK |
| --- | --- | --- | --- | --- |
| Reviving stage | SPAD | 0.2823* | 0.146 | 0.1683 |
|  | LAI | 0.2097* | 0.1147 | 0.1426 |
| Jointing stage | SPAD | 0.5007** | 0.1598 | 0.2063* |
|  | LAI | 0.5412** | 0.1737 | 0.2587* |
| Filling stage | SPAD | 0.5168** | 0.1956 | 0.2136* |
|  | LAI | 0.5635** | 0.2025* | 0.2645* |

Note: \*\*P<0.01; \*P<0.05. The same below.

With the increase or decrease of AN content in the soil, the values of SPAD and LAI also increased or decreased correspondingly during the period of reviving stage to filling stage. With the increase of AP content in the soil, the values of LAI also increased correspondingly at filling stage, while the change of SPAD at filling stage and SPAD, LAI at the other three stages were not obvious. It showed that AP had different effects on wheat growth at different growth stages. The values of SPAD and LAI were significantly correlated with the AK content in the soil during jointing stage to filling stage, and the correlation coefficients between this two growth stages were not significantly different. It indicated that AK played an important role during the late growth stage of wheat, and had a steady effect on wheat growth compared with AN, AP.

There were the correlation coefficients between the content of soil nutrient at the current stage and the index of wheat growth at the next growth stage (Table 7). It could be seen that there was also a good correlation between the content of soil nutrient at the current stage and the index of wheat growth at the next growth stage. Among them, there was extremely significant correlation between AN (before sowing) and SPAD (reviving stage), AN (jointing stage) and SPAD, LAI (filling stage). There was significant correlation between AN (before sowing) and LAI (reviving stage), AK (before sowing) and SPAD (reviving stage), AK (jointing stage) and SPAD, LAI (filling stage).

**Table 7:**
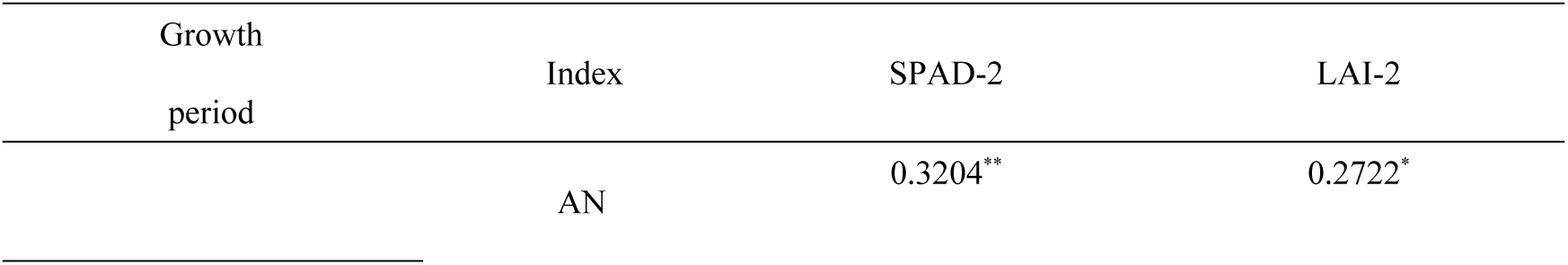
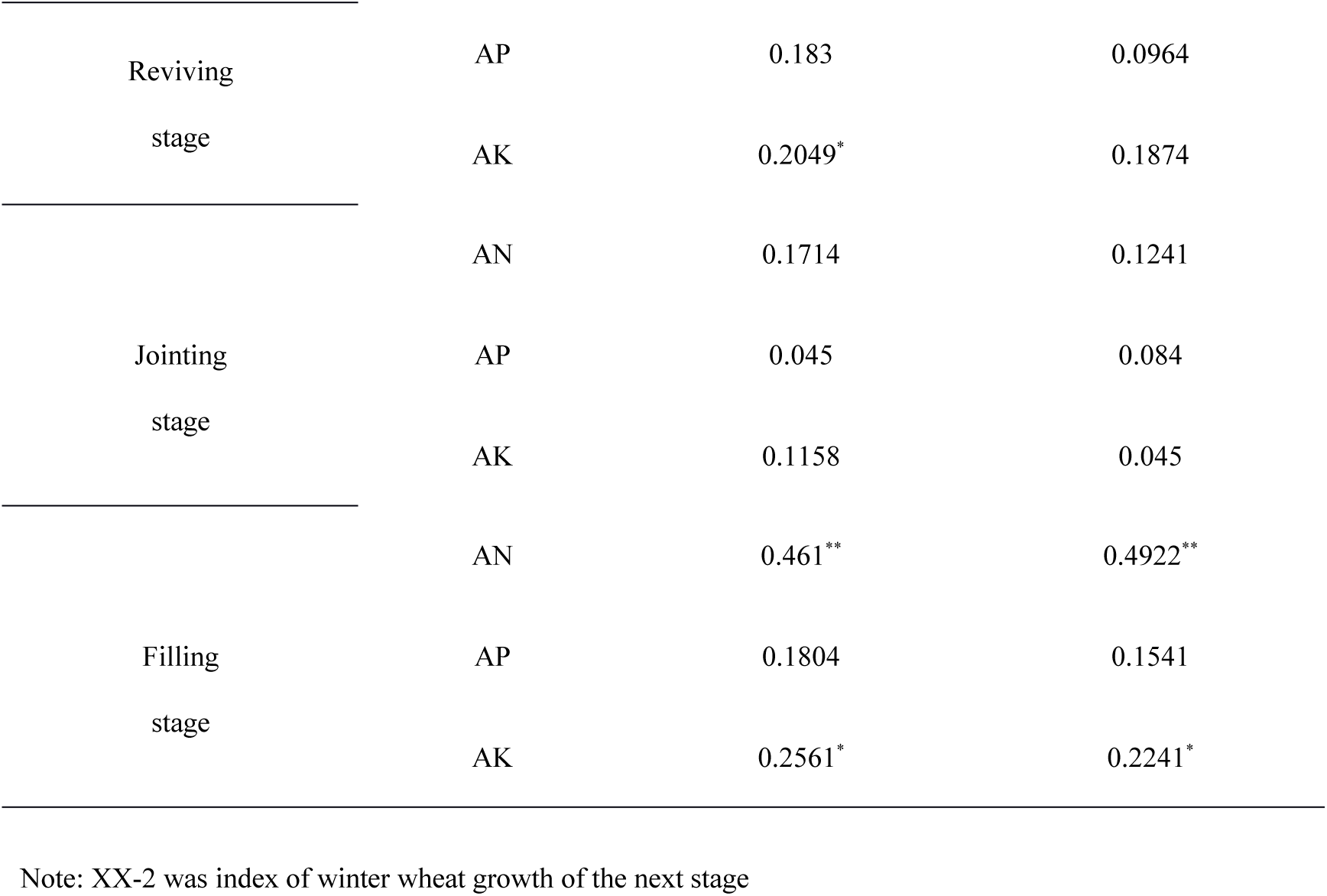
Correlation coefficients between soil nutrients and wheat fertility index of the next growth period.

In comparison table 6 and table 7, the correlation coefficients between AN, AP and AK (before sowing) and the value of SPAD, LAI (reviving stage) was significantly higher than the correlation coefficients between AN, AP and AK (reviving stage) and the value of SPAD, LAI (reviving stage). This indicated that the nutrients of before sowing had more obvious effect on the growth of wheat at reviving stage than the nutrients of reviving stage. The correlation coefficients between AN, AP and AK (reviving stage) and the value of SPAD, LAI (jointing stage) was significantly lower than the correlation coefficients between AN, AP and AK (jointing stage) and the value of SPAD, LAI (jointing stage). This indicated that the growth of wheat at jointing stage was mainly affected by the nutrient of the current stage, which was related to the jointing stage being a peak period of nutrient uptake by wheat. This was mainly related to the reason of jointing stage was a peak period of soil nutrient absorption by wheat. The correlation coefficients between AN, AP and AK (jointing stage) and the value of SPAD, LAI (filling stage) was significantly lower than the correlation coefficients between AN, AP and AK (filling stage) and the value of SPAD, LAI (filling stage), but it still reached significant correlation. This indicated that the growth of wheat at the filling stage was both affected by soil nutrient of jointing stage and filling stage

In general, the growth status of wheat was closely related to the nutrients of soil. AN had the highest correlation with wheat growth status, followed by AK and AP lowest. At reviving stage, the effect of nutrients of before sowing on the growth of wheat was slightly higher than that of reviving stage. At jointing stage, the effect of nutrients of jointing stage on the growth of wheat was much higher than that of reviving stage. At filling stage, the status of wheat growth was affected by the nutrients of jointing stage and filling stage.

## Discussion

(1) The method of experimental design has the advantages of easy access to data, low cost and small interference, but it is different from the real field environment and management mode, and the conclusion has great limitations in the application of agricultural extension. Based on this, this research studied the soil nutrients and winter wheat under the traditional tillage model, and the conclusion was more practical, and could be better suitable for winter wheat planting area in northern China.

(2) Huantai Country was located in Huanghuai winter wheat area, the main wheat production area of China, with superior planting conditions. It was representative to study the variation of soil nutrients and their correlation with crop growth during the growth period of winter wheat here. The minimum spatial autocorrelation distance of soil nutrients in each growth stages was 88.2m, which exceeded the sampling interval (60m), and indicated the sampling interval was reasonable. In this study, the dynamic changes of soil nutrient content during winter wheat growth period were generally consistent with the research on Jiangyan City, Jiangsu Province, China [28].

(3) It was found that soil nutrients showed moderate variation during the growth stage of winter wheat, generally showed that AN was slightly larger than AP, but much larger than AK. The reason was mainly related to fertilization. The high variability of AN was due to the reason of wheat has a large demand for nitrogen and farmers pay great attention to the use of urea. The high variability of AP was related to the chemical behavior of P elements in the soil [29], its mobility was weak and its utilization rate was low, resulting in a large amount of fertilizer residue in the soil and uneven distribution, which was also confirmed by the maximum variation of AP before sowing in this research. The soil parent material in the study area was rich in potassium, which has surplus for the supply of the growth of wheat. Therefore, the influence of field management on AK was small, so the variability was low.

(4) It was found that single C_0_/(C_0_+C) could not be stable and effective in describing the structure of the variable when C_0_/(C_0_+C) was large in a small area [30]. Therefore, the method of mutual verification of C_0_/(C_0_+C), fractal dimension and Moran ’s I was adopted in this research. The results showed that the structure of soil nutrients weakened from before sowing to reviving stage, jointing stage to filling stage, while tended to strengthen from reviving stage to jointing stage. This may be caused by the field management and the nutrient absorption of wheat growth. Especially, the urea supplementation at reviving stage and irrigation at filling stage were the main reason for the obvious decrease of soil nutrient autocorrelation. Nutrient uptake of winter wheat reached its peak at jointing stage and soil nutrient distribution tended to be balanced, then structural enhanced. It was also found that there were significant changes of soil nutrient autocorrelation during the growth period of tobacco [31]. Wheat, one of the most important food crops in the world, was of great significance to study the effects of wheat growth on soil nutrients systematically, However, the results in this field still remain to be enriched.

(5) The correlation coefficient between AN and SPAD, LAI was the highest at all growth stages of winter wheat, followed by AK and AP was the lowest. This may be related to the difference in the amount of demand for different nutrients during the growth stage of winter wheat. It was found that the ratio of wheat to nitrogen, phosphorus and potassium absorption was 3.1:1:3.14 in the brown soil area of western Henan province, China [32], similar to the planting conditions of this research area. It was basically consistent with the conclusions of this study. The correlation between the soil nutrient and the growth index of the next growth period showed that the growth of the winter wheat of reviving stage was mainly influenced by the soil nutrients of the previous period (before sowing), indicating that there was a time delay from fertilization to fertilizer affects crop growth. The effect of soil nutrients at jointing stage on the growth of wheat at jointing stage was much higher than that of soil nutrient jointing stage. This was related to the reason of high demand for fertilizer at jointing stage of wheat [33]. Therefore, ensuring the supply of fertilizer at jointing stage was of great significance. At filling stage, the effect of nutrients of previous stage (jointing stage) on the growth of winter wheat was still obvious. Because the traditional period of urea supplementation was generally chosen at reviving stage in the area of northern China, in view of the importance of the jointing period to the growth of winter wheat, the time of the fertilizer supplementation should be properly moved to the jointing stage to ensure the nutrient supply in the later period of the wheat growth. The spatial and temporal distribution of more biological indicators of winter wheat need to be further studied.

## Conclusion

(1) During the growth stages of winter wheat, three nutrients all showed the characteristics of decreased-increased-decreased in content and moderate variability. The reviving stage was the turning point of the content of AN, AK and AP was the jointing stage, its content increased before the stage, while decreased continuously after the stage. The coefficient of variation of AN was the largest, AP was the second, and AK was the smallest.

(2) The best fitting model of the three nutrients before sowing were the same, AP and AK were same at reviving stage, the three were different at jointing stage, AN and AP were the same at filling stage, difference of nutrient variation first increased and then decreased. The three nutrients all showed medium to strong spatial correlation during the growth period. The three nutrients were the weakest spatial autocorrelation at reviving stage; the before sowing of AN was the strongest, while the AP and AK were the strongest at jointing stage. The autocorrelation of soil nutrient decreased from before sowing to reviving stage, jointing stage to filling stage; and increased from reviving stage to jointing stage.

(3) Kriging interpolation analysis showed that with the growth of winter wheat, the grade of high-content level of AN gradually decreased and spatial distribution increased continuously; the grade of high-content level of AP decreased slightly, the change of spatial distribution continuity was not obvious; the grade of high-content level of AK decreased significantly and spatial distribution continuity decreased continuously. The content of each nutrient generally experienced a process of increasing first and then decreasing gradually, the number of nutrient grades, especially the number of high-content grades, decreased gradually.

(4) Pearson correlation analysis showed that the correlation coefficients between soil nutrients and the index of wheat growth gradually increased with the development of winter wheat. AN had the highest correlation with wheat growth status, followed by AK and AP lowest. At reviving stage, the effect of nutrients of before sowing on the growth of wheat was slightly higher than that of reviving stage. At jointing stage, the effect of nutrients of jointing stage on the growth of wheat was much higher than that of reviving stage. At filling stage, the status of wheat growth was both affected by the nutrients of jointing stage and filling stage. Fertilization of winter wheat should pay attention to applying basal fertilizer and the time of fertilizer supplementation should be properly delayed around to the jointing stage.

In this research, the temporal and spatial variation characteristics of soil nutrients and the response of wheat growth in the whole growth period of Winter Wheat under the traditional tillage model were well understood, which could provide a scientific basis for the effective guidance of wheat production and the popularization of precision fertilization technology in the broad winter wheat area of northern China.

## Acknowledgements

We thank the agriculturist in the Agricultural Bureau of Huantai County for their help on the field work. We also thank the language expert of Elsevier language editing service for help to improving the language quality of this manuscript, and the anonymous reviewers for reviewing this manuscript and putting up comments.

## Author Contributions

Data curation: Baowei Su, Chao Dong.

Investigation: Baowei Su, Chao Dong.

Project administration: Baowei Su, Gengxing Zhao.

Software: Baowei Su, Chao Dong.

Methodology: Baowei Su, Gengxing Zhao, Chao Dong.

Validation: Baowei Su.

Writing - original draft: Baowei Su, Gengxing Zhao.

Writing - review & editing: Baowei Su, Gengxing Zhao.

